# G-quadruplexe as a structural modulator of Intron Retention upon viral infection

**DOI:** 10.1101/2023.04.12.536615

**Authors:** Pauline Lejault, Michel-Pierre Terrier, Anaïs Vannutelli, François Bolduc, Carolin Brand, Martin Bisaillon Jean Pierre Perreault

## Abstract

Amongst the wide array of alternative splicing events (ASE), the intrinsic mechanisms of intron retention have remained elusive. This particular type of ASE has long been characterized as an artifact, but recent studies have shown its implication in numerous diseases. It has also been revealed that numerous viruses choose to disrupt alternative splicing to escape cellular immune response and further their proliferation. The main focus of this study was to investigate the G-quadruplex role in Alternative Splicing Events (ASEs) that occur following Flavivirus infections. After having demonstrated that G-quadruplexes structures are mainly formed in Intron Retained Transcripts by RNA-seq, our attention turned toward the ULK3 gene, coding for a serine/threonine kinase regulating autophagy, an essential mechanism in the cellular response to stress and even pathogen infections. In this study, we revealed the presence of a G-quadruplex in the first intron of the ULK3 gene near the 3 ’ splice site. Furthermore, we assayed the formation and stability of this G-quadruplex in vitro and showed that its formation affects IR, as demonstrated by comparisons between wild-type and mutant transfected mini-genes. Finally, we identified the specific RNA-binding protein signature for this G-quadruplex, thereby uncovering the novel role of G-quadruplexes in Alternative Splicing.

## Introduction

First characterized in 1977, the phenomenon of alternative splicing (AS) is an essential cellular mechanism from which genes can produce multiple protein isoforms with distinctive functions, different cellular localization, or marked for degradation^1^. This mechanism allows the relatively small number of human genes to express a great variety of mRNA and is one of the hallmarks of our proteomic complexity ^2,3^. As can be easily disrupted by cellular stress or different diseases including cancer and Amyotrophic Lateral Sclerosis ^4,5^. Incidentally, studying the wide array of alternative splicing events (ASEs) in genes remains an interesting track to decipher the hidden mechanisms of alternative splicing. Amongst those five main ASEs such as exon skipping or alternate 5′ or 3′ splicing, intron retention remains the most understudied ^6^. A growing number of articles estimated that more ASEs are induced by adverse stimuli that occur to confer stress tolerance^7^. In eucaryotes, various genotoxic compounds, physical stressors such as heat and light (plant), or viruses can be used to modulate the generation of ASEs ^7–9^. We recently reported how orthoreoviruses can induce ASEs by targeting the core splicing machinery by reducing the availability of the host U5 snRNP proteins^10^. These encouraging discoveries have not only shown the complex relationship between host and virus in viral infections but also the need to further our understanding of the underlying mechanisms of AS.

Beyond these elements of external stress, removal intronic sequences process from the pre-messenger RNA necessitates three fundamental signals such as the *branch point*, the *3′ Splicing Site*, and the *5′ Splicing Site* ^3^. However, *In silico* studies have demonstrated that only half of these signals determining the exon/intron boundaries are present in transcripts, thus suggesting that another signal or structure might be present and involved in the recognition of exon/intron splicing sites ^3,11^. *Georgakopoulos-Soares et al*, based on RNA-seq analysis, concluded that G/C-rich sequences near the splicing junction appear to promote the formation of G-quadruplexes ^12^. Additionally, retained introns tend to be short and have a higher G/C concentration, which highlights the significance of this structure in ASEs^13^.

These secondary structures have a high potential of formation with more than a million of these predicted to form a G-quadruplex^14^. Depending on their localization on pre-mRNA, RNA G-quadruplexes (rG4) have been shown to modulate critical steps in AS. Still, the mechanism remains controversial: rG4s have the potential to act either as roadblocks (repressors) for different elements of the splicing machinery (they may conceal cis-regulatory elements) or as enhancers and serve as binding sites for RNA-binding proteins (RBPs) ^15^. For instance, a rG4 located in intronic region 6 of the human telomerase transcript (hTERT) has been suggested to regulate hTERT splicing efficiency as a splicing attenuator which facilitates exon skipping and results in the production of inactive hTERT protein^16,17^. On the other hand, rG4 located in intron 3 of the TP53 pre-mRNA facilitates the splicing of the adjacent intron ^18^. Moreover, several proteins that regulate AS have been identified to bind G4 motifs through proteome-wide pull-down experiments^19^. Some hnRNPs and splicing factors exhibit high affinity to G4 motifs, for instance with hnRNPK/U ribonucleoproteins positively regulating exon inclusion and FMRP enhancing the splicing of its own transcript by G4 binding^12,20^.

To summarize, the presence of rG4 structures can significantly impact alternative splicing processes in positive or negative ways. To reconcile conflicting findings and make it easier to study ASEs events, we explored how rG4s can affect AS, particularly in response to viral infection. We used RNA-seq data to predict rG4 formation in ASEs following three different flavivirus infections, including ZIKA, Yellow Fever, and Kunjin. Amongst the 2,253 ASEs detected, Intron Retained Transcripts (IRTs) were found to be the most pG4-rich events. Our research subsequently centered on ULK3 IRTs, which were consistently identified in all three virus infection scenarios. We also confirmed the existence of rG4 and its influence on IR by minigene. Finally, we employed rG4 ULK3 to capture RNA-binding proteins (RBPs) and investigate the potential mechanism by which rG4 might contribute to IR.

### Acquiring datasets for precise detection of Alternative Splicing Events and pG4 quantification

For several decades, the modulation of the number of ASEs in the host cell has been observed in numerous viral infections caused by different types of viruses, including double-stranded DNA viruses (e.g: Herpes simplex virus,^21^ Epstein-Barr virus ^22^) and RNA viruses (e.g: double-stranded such as Reovirus^10^ or single-stranded such as Sindbis virus ^23^). To investigate the landscape of G4s present in ASEs *in vivo*, we performed viral infection with three flaviviruses (e.g Zika (ZIKV), Yellow Fever (YFV), and Kunjin (KUNV)) as a source of stress for host cell transcriptome (glioblastoma cell line U87). Flaviviruses are a family of positive-strand RNA viruses previously known to significantly affect pre-mRNA splicing and alter the host cell ‘s transcriptome ^24–26^.

Once cells were infected with Zika, Yellow Fever, and Kunjin (MOI=5), RNA was extracted and purified after 24 hours to identify ASEs using RNA-sequencing (Figure 1.A). To ensure the accuracy of ASE identification a data processing pipeline was established. The pipeline comprised four key stages, beginning with a cleaning step to remove low-quality adapters and reduce read length using trimmomatic^27^. The read mapping to the hg38 reference human genome was then conducted using STAR^28^, followed by differential analysis of AEs through comparison of infected cells versus control conditions using rMATS^29^. The expected increase in AEs was observed with 222,391 AEs detected involving KUN, YFV, and KUNV. In order to ensure the reliable detection of AEs, viral infection was performed in triplicate and the identified AEs were filtered based on three scores: ΔPSI > 0.1, Pvalue <0.05, and a FDR<0.05. A total of 2253 ASEs with high confidence were detected, which corresponded to the top 1% of all AEs (See figure SUPP). The ultimate stage of the process consisted of using G4RNAScreener to determine whether the type of AE was related to the density of potential G4 sequences ^30^. This tool assessed the number of predicted G4 motifs identified, using three stringent scoring criteria, in the short window of 100nt upstream and downstream to the splicing junction of AEs. We observed that intron retention events are the most enriched in pG4 sequences, with 20% of pG4 motifs present in all three virus conditions (see figure 1B). Although rG4s may play an important role in various AEs our finding confirms the previously suggested implication of rG4s in IRTs, likely due to the high G/C percentage of these events^13^. Moreover, one of the most well-known examples of intron retention (IR) events that involve the formation of rG4s is located in intron 1 of C9ORF72, which is a known cause of amyotrophic lateral sclerosis (ALS) and the sequestration of splicing factors, such as hnRNPH ^4,6^. Based on these findings, we further analyzed the common intron retention events among the three viral infections that contained at least one pG4 motif. After evaluating both candidates, we have concluded that the ULK3 transcript is the best candidate for studying rG4 formation in the context of IRTs (as shown in Figure 1C) The decision to opt for ULK3 instead of MAPK12 was based on the fact that the possible rG4 location in ULK3 does not fall within a splice site, unlike MAPK12. Therefore, any modification required to study rG4 in this region may disrupt the recruitment of splicing machinery (Figure 2A). Moreover, ULK3 had an exceedingly high G4RNA score (0.9968).

**Figure 1.**
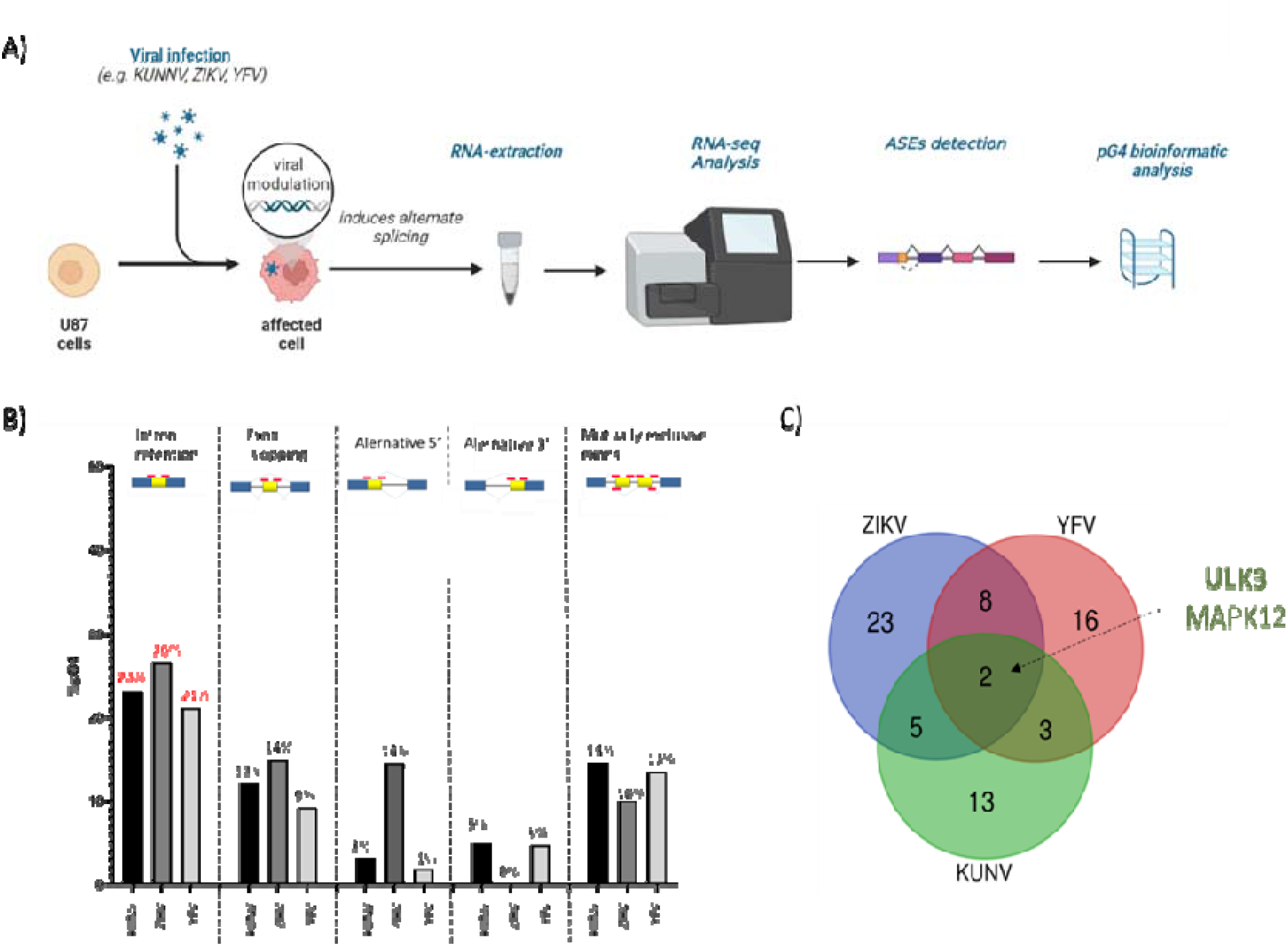
Identification of pG4 in alternative splicing events. A) Schematic representation of the experimental workflow used to identify ASEs and pG4 B) The results obtained, indicating the percentage of pG4 found in AEs detected in three different viral infections MOI=5 (Kunjin KUNV, Zika ZIKV, and Yellow Fever YFV). C) The Venn diagram represents intron retention events that have at least one pG4 in common across the three flavivirus Infections.

**Figure 2.**
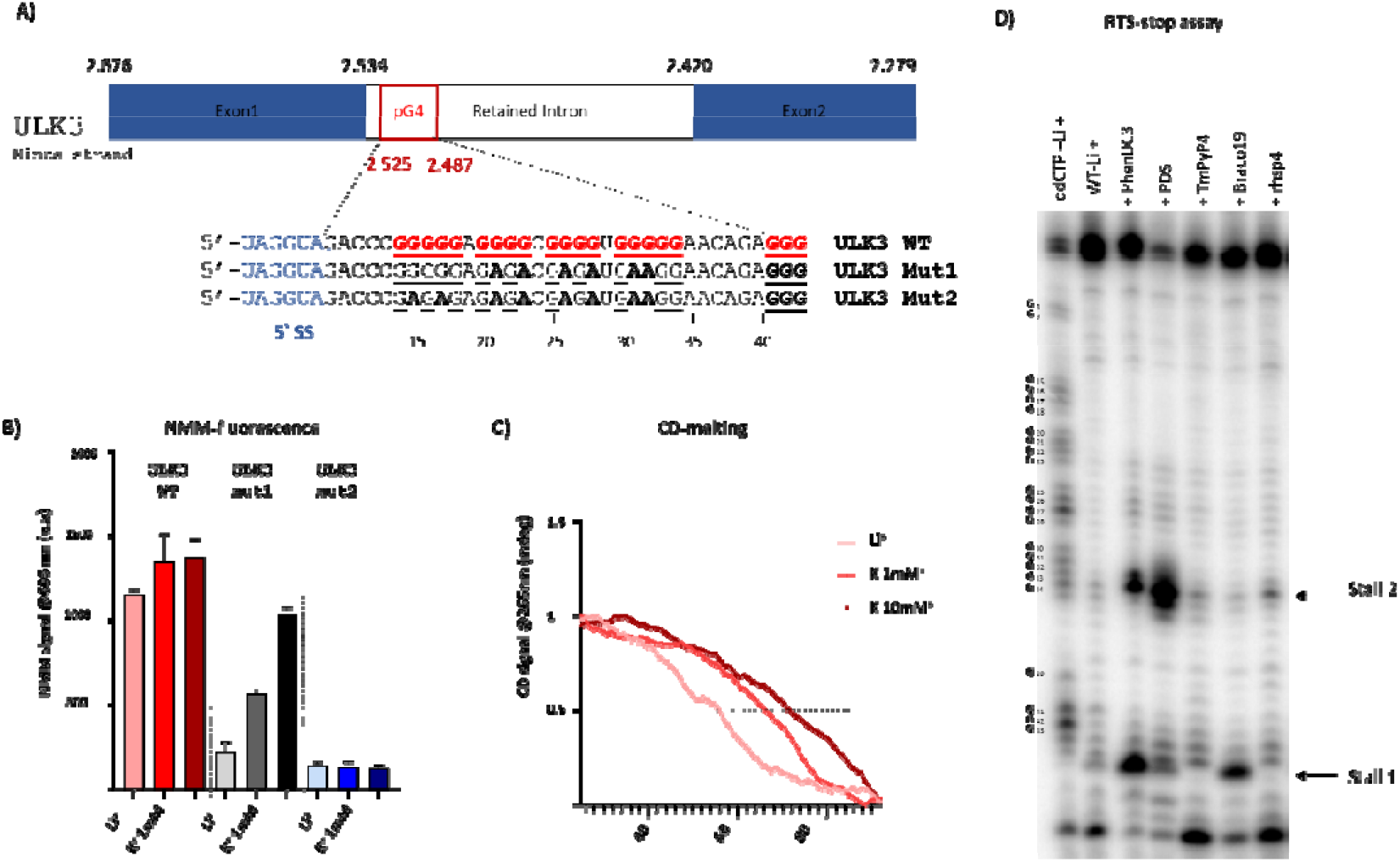
Biophysical characterization of RNA secondary structure formation. A) Description of Intron retention 1 in ULK3 and the localization of G-rich sequence in the wild type mRNA and the relative mutants. B) Results of NMM fluorescence assay with ULK3 wt,mut1,mut2 in cacodylate buffer with Li^+^, 1mM K^+^ and 10mM K^+^). C) Spectra of CD melting of ULK3 WT in cacodylate buffer with Li^+^, 1mM K^+^ and 10mM K^+^ and D) RTS-stop assay results, with ULK3 WT in presence of PhenDC3, PDS, Tmpyp4, Braco19 and RHPS4 (10mol.equiv).

### rG4 structure affects the splicing of ULK3 mRNA

Unc-51 like kinase 3 (ULK3) gene is known to be involved in the crucial autophagy pathway, which is responsible for breaking down and recycling intracellular components. ULK3 plays a significant role in initiating autophagy by contributing to the formation of the ULK1-ATG13-FIP200 complex^31^. Moreover, ULK3 kinase helps regulate the activity and positioning of other autophagy-related proteins such as Beclin-1 and LC3^32^. Among the transcript variants of ULK3, the ASE was identified in an intriguing coding transcript (ENSC00000140474), where a guanine-rich sequence (ULK3-WT) was found positioned between Exon 1 and Exon 2, located just 5 nucleotides away from the 5′ spliced site (as shown in Figure 2-A). Although G4RNAscreener is a powerful tool to identify rG4s, it’s important to validate the formation of this specific structure in vitro. Therefore, a set of well-established in vitro experiments (including NMM fluorescence assay, CD-melting, and RTS-Stop assay) were performed to confirm its formation^33–35^.

Two mutant variants, ULK3-mut1 and ULK3-mut2, were obtained to study the impact of the pG4 structure due to its proximity to the splice site. In ULK3-mut1, the first 5 guanines remained unchanged, while in ULK3-mut2, all stretches of guanines were substituted with adenines. (Figure 2A). To evaluate the formation of rG4 structures in different constructs, N-methylmesoporphyrin IX was added as a turn-on probe (Figure 2B)^36^. This probe enables the detection of parallel G4 structures, with a signal increase at 606 nm observed with ULK3 WT in the presence of K^+^ (at 1mM and 10mM, which are known to strongly promote G4 formation) and Li^+^ (which is less favorable for G4 formation). The impressive thermal stability of ULK3-WT confirms its formation, with Tm values of 54°C, 65°C, and 71°C in Cacodylate buffer containing 10mM Li+, 1mM K+, and 10mM K+, respectively (Figure 2C). Interestingly, ULK3-Mut2 appears to form an rG4 structure that shows increasing fluorescence as the saline concentration of K^+^ increases (u.a= 1030.5 k^+^10mM). Despite six nucleotides being interchanged, it seems that an rG4 structure can still form a 2-Gquartets G4 in vitro, unlike in ULK3-mut2 where fluorescence remains low (<150 u.a).

To gain a better understanding of the precise location of the rG4 structure and its potential stabilization with G4-ligands, RT-stall assays were performed^35^. As previously described, this technique detects reverse transcriptase stalling (RTS) events that indicate the formation of rG4 by leveraging the specific regulation of rG4-cation and G4-ligand interactions during reverse transcription (Figure 2D and SUPP). The used of ligands, specifically PhenDC3, PDS, and BRACO19, seems to impact the formation of rG4 within ULK3 WT as depicted in Figure 2D. Two different stop sites were identified under these circumstances, associated with either the binding of the first G stretch (G_45_) or the following (G_34_). This observation suggests the existence of two distinct rG4 structures, probably comprising three and four G-quartets. While in the case of ULK3-mut1, rG4 formation appears to occur only when stabilized by PhenDC3 and PDS, no stop was observed in K^+^ (Figure supp A,B). and as expected, any stall have been observed with the ULK3-mut2 control.

### Impact of rG4 on ULK3 splicing *via* minigene constructs and its protein partners

Upon validating the formation of rG4 structures in ULK3, we proceeded to create three minigenes to provide direct evidence of the impact of rG4s on intron retention. Additionally, we aimed to identify the RNA-binding proteins that mediate this regulatory mechanism, following a workflow illustrated in Figure 3A.

**Figure 3.**
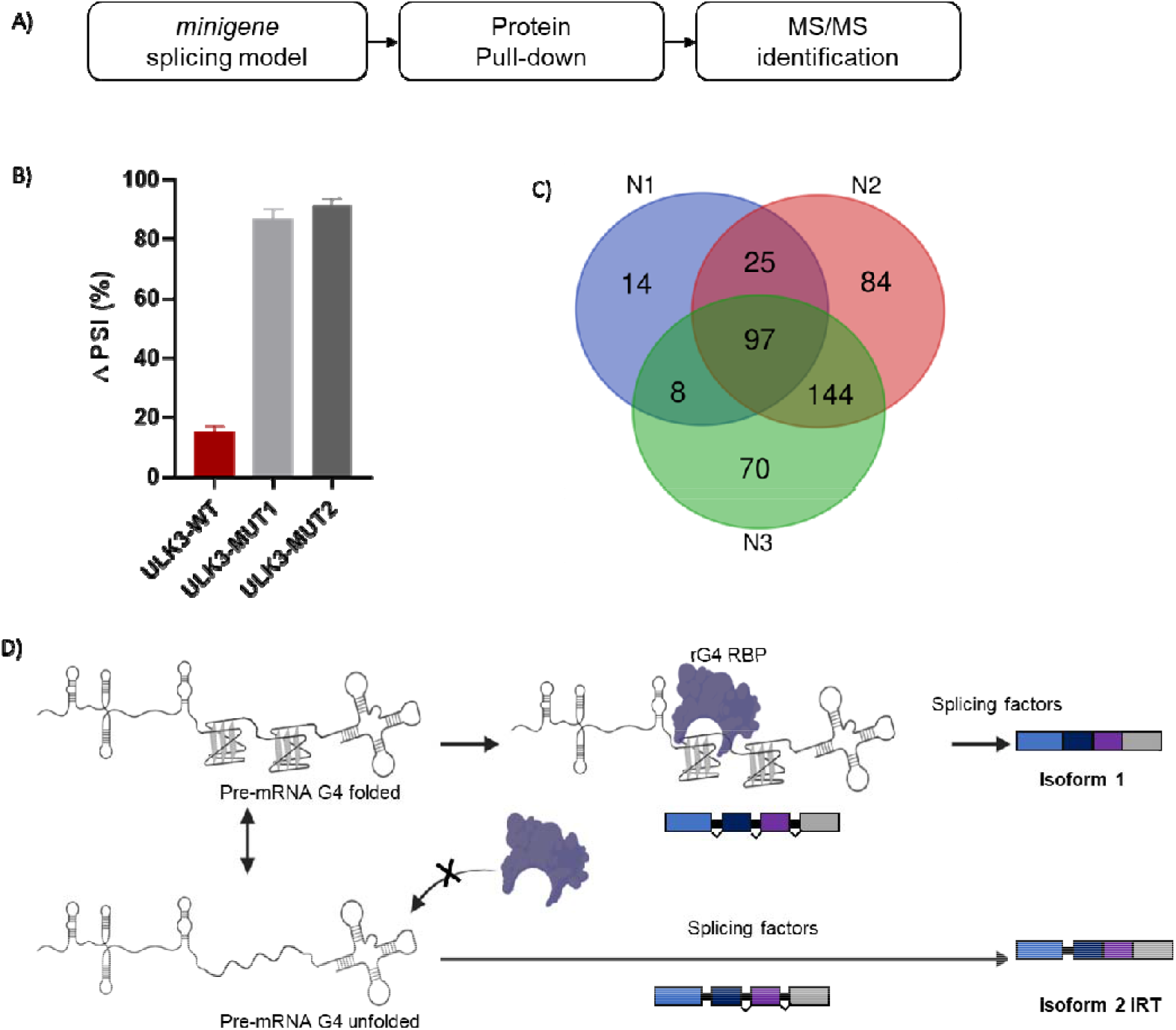
A) Workflow of viral infection and RNA-seq data treatment B) RT-PCR results of ULK3-WT, ULK3-MUT1 and ULK3-MUT2 constructs DPSI = (long form)/(long form+ short form). C) Venn diagramm of RBPs enriched in presence of G4 ULK3 by LC MS/MS. D) Schematic representation of relationship between G4s and intron retention of ULK3:G4s structures can modulate the splicing via recruitment of RBPs.

For the ULK3 minigenes, we inserted either the wild-type (ULK3-WT) or mutated G4 motifs (ULK3 mu1 and ULK3 mut2) between two native exons (Exon 1 and Exon 2).After 24h of this transitory transfection, RNAs was purified and analyzed by RT-PCR. The findings are remarkable as they reveal a considerable decrease (15%) in the percent spliced in (ΔPSI) of intron retention when the rG4 is present. In contrast, when the rG4 wasn’t fold, in mut1 and mut2, the long form RNAs were predominantly produced, leading to 87% and 91% intron retention, respectively. To validate these findings, the experiment was replicated in other cell lines (HEK293T) using a reduced number of transfected minigenes (250ug to 3000ug) to prevent the saturation of splicing machinery. (Figure Supp). In summary, the rG4 structure plays a critical role in facilitating proper alternative splicing, as its absence results in the retention of intron 1. While there are only a few reported instances of rG4 aiding splicing, the targeting of the INS gene with ASOs highlights the significance of rG4 in therapeutic target RIs.

To identify RBPs capable of binding to the rG4 structure and regulating ULK3 differentially, a protein pull-down strategy was employed. A chimeric RNA composed by 180 nucleotides RNA containing intron 1 with the wild-type rG4 fused to four repeats of the S1m aptamer, which is a modified RNA aptamer that binds to streptavidin have been used. The chimeric RNA was immobilized on streptavidin-coated sepharose beads. Next, total protein extracts from U87 cells were incubated with the chimeric RNA. Non-specific interactions were washed away, and the bound RBPs were subjected to trypsin digestion. The resulting peptides were analyzed using LC-MS/MS spectrometry to determine which proteins were enriched in the wild-type rG4 in K^+^ samples (rG4 is fold) compared to Li^+^ (rG4 isnt fold) samples. The experiment was performed in triplicate, and 97 potential candidates that showed a fold enrichment of >2.0 were identified (Figure 3C).

Among these candidate, several RBPs were already known to be rG4 helicases or chaperones, such as DDX21^37^, DDX5^38^, DDX3X^39^, DDX17^39^, nucleolin^40,41^, which confirmed the specific identification of G4RBPs. Surprisingly, 10% of the identified proteins were known as splicing actors, including members of the Heterogeneous Nuclear Ribonucleoprotein family (HNRNP), such as HNRNPD, HNRNPA2/B1, HNRNPU, HNRNPL, HNRNPR, HNRNPA, HNRNPK, as well as splicing factors i.e: USP39, PRPF38B, and SF3B3. Moreover, HNRNPs are RNA-binding proteins that play crucial roles in RNA processing, including splicing, transport, and stability. Some of them were also found to be capable of recognizing rG4s,, including FUS^42^, HNRNPU^12^, HNRNPK^12^, and HNRNPA2/B1^43^ (Figure Supp). Interestingly HNRNPD, HNRNPL, and HNRNPR, were not previously recognized as G4 binders, despite showing significant enrichments of 6-fold, 5-fold, and 4-fold, with ULK3 construct respectively. These RNA-binding proteins contain several RGG (arginine-glycine-glycine) motifs, which are known to be enriched in G4 RBPs.

Moreover USP39, PRPF38B, and SF3B3 have not been previously reported to be involved in rG4 recognition. USP39 is a component of the spliceosome^44^, while PRPF38B is involved in pre-mRNA splicing^45^, and SF3B3 is a core component of the U2 snRNP complex, which is also involved in pre-mRNA splicing ^46^.

The discovery of new RNA-binding proteins, including HNRNPs and splicing factors, that recognize G-quadruplex structures expands our knowledge of the intricate mechanisms that regulate RNA processing and provides new insights into the role of G4 structures in splicing regulation. Moreover, intron retention, has been implicated in a wide range of diseases, such as cancer, muscular dystrophies, and neurodegenerative disorders. By identifying the G4 RBPs involved in intron retention, we gain a better understanding of the regulation and mechanisms underlying this process and open up new avenues for developing targeted therapeutic interventions for diseases caused by RNA splicing mis-regulation.

## Conclusions

In conclusion, this study sheds light on the previously elusive mechanisms of intron retention, which has been shown to play a significant role in various diseases. By investigating the role of G-quadruplexes in Alternative Splicing Events (ASEs) following Flavivirus infections, the study revealed the presence of a G-quadruplex in the first intron of the ULK3 gene, which regulates autophagy, a critical mechanism in the cellular response to stress and pathogen infections. The study also demonstrated the formation and stability of the G-quadruplex near the splice junction can impact its intron retention.

The specific RNA-binding protein signature for this G-quadruplex was identified, highlighting the novel role of HRNPs proteins (HNRNPD, HNRNPL,HNRNPR) and also three splicing factors (USP39, PRPF38B, and SF3B3) in the recognition of rG4 and its importance in Alternative splicing. Overall, these findings provide valuable insights into the complex interplay between G-quadruplexes, ASEs, and viral infections, with potential implications for the development of novel therapeutic strategies

## Experimental part

### Cell culture

Human Kidney Embryo 293T (HEK293T) and Human Glioblastoma Astrocytoma (U87-MG) cells lines were cultured in Dulbecco’s modified eagle medium (DMEM) supplemented with 10% FBS for both, plus 1mM sodium pyruvate (U87 only). All cells were cultured at 37°C in a humidified 5% CO2 incubator.

### Viral infection

Kunjin virus (strain FLSDX, GenBank AY274504.1), Zika virus (strain PRVABC59, GenBank KU501215.1), and Yellow Fever virus (strain 17D, GenBank X03700.1) were used for viral infection. To perform infections, 50% of confluent U87 in T150 cells were either infected at a multiplicity of infection (MOI) of 5 or treated with cell-conditioned media (n=3 for each virus and control). 24h post-infection, cells were harvested and homogenized using QIAshredder (Qiagen). RNA was isolated using the RNeasy kit (Qiagen) and stabilized for storage using an RNA transport kit (OMEGA bio-tek, R0527). Then, PolyA-RNA was purified from 5µg total RNA using NEB magnetic mRNA Isolation Kit (NEB, #S1550S) and eluted into a final volume of 25µl. Sequencing libraries were prepared using 9µl of isolated mRNA and ScriptSeq RNA-Seq Library Preparation Kit (Epicentre, SSV21124). Paired-end 100 bp sequencing was performed on a HiSeq 4000 system (Illumina) at McGill University and Génome Québec Innovation Centre, obtaining between 34 and 72 million reads per sample.

### Bioinformatics analysis

The bioinformatic pipeline consists in 4 steps to identify the ASEs (i.e: trimmomatic, star and rMats) and the pG4s positions (i.e: G4RNA):

#### Trimommatic

Reads were trimmed using Trimmomatic 0.32 to remove adapter sequences as well as nucleotides with low quality scores (TRAILING: 30) ^27^. The quality of output reads was verified using FastQC 0.11.5. <u>Star :</u> The data were aligned to the reference genome (hg38 + viral genomes) using STAR 2.5.1b with default parameters ^28^. <u>rMATS</u> : Alternative splicing events analysis was performed using <u>rMAT</u>S 3.2.5 with options -len 93 -a 3 -t paired -analysis U ^29^. ΔPSI (percent spliced in) values for each alternative splicing event were calculated by subtracting the PSI value in infected cells from the PSI value in non-infected cells. Therefore, a positive ΔPSI value indicates an increase of the short form upon infection, whereas a negative ΔPSI value indicates an increase of the long form upon infection. Finally, to be considered as ASEs significant, 3 filters was necessary : DPSI > 0.1, P value < 0.05,FDR <0.05 (See figure S1). And finally,G4RNAscreener : It was launched on the full unspliced gene sequences identified as ASEs with 100-nt located upstream and 100-nt downstream of the splicing junction. The screening used a sliding window approach, with a window length of 60 nt and a step of 10 nt (i.e. the recommended default parameters).The threshold score limits for pG4 detection were set to 4.5 for the cG/cC score, 0.9 for G4H and 0.5 for G4NN (as recommended for stringent detection) ^30^.

### Transfection

Cells were seeded in six-well plates (200x 104 cells/well u87 ; 50 × 104 cells/well HEK293T) and transient transfections were carried out 24h later using the Lipofectamine ™ 2000 reagent (invitrogen) according to the manufacturer ‘s protocol. Initially, 30o0ug of plasmide was used in U87 experiments. Then, a concentration gradient was also tested in HEK293T (250ug, 500ug, 1500ug and 3000ug).

### Molecular cloning

The PcIDT plasmids containing either Exon1-Intron1-Exon2 of ULK3 WT sequence were purchased from IDT. The regions of interest were then co-digested with EcoRI-HF and Hind III-HF, and ligated into the mammalian expression vector pcDNA3.1. The resulting plasmids were amplified in the Escherichia coli DH5α strain (see Table S1 for the full sequences). For RBP experiments, the plasmid pMARQ_S1mX4 containing four repeats of the S1m aptamer was constructed as previously described47. The plasmids pMARQ_ULK3_S1m4X was generated by PCR-amplifying the region containing Exon1-Intron1-Exon2 of ULK3 WT from the PcDNA3.1 plasmids, digesting it, and cloning it into the pMARQ_S1mX4 construction. All of resulting plasmids were amplified in the Escherichia coli DH5α strain and then, plasmid purifications were performed according to the protocol provided with the PureLink HiPure Plasmid Midiprep kit (Invitrogen). All constructions were confirmed by PCR-colony and sequenced at the DNA sequencing platform of Laval University.

### Biophysical characterization of G4s

All oligonucleotides used here were purchased from Integrated DNA technology (IDT) and stored at -20°C at 100 to 1000 µM stock solutions in deionized water. The actual concentration of these stock solutions was determined through a nanodrop measurement at 260 nm.

#### Circular Dichroism spectra and CD-melting

CD were recorded on a JASCO J-810 spectropolarimeter in a 1 mm path-length quartz semi-micro cuvette. CD spectra were recorded over a range of 210-350 nm (bandwidth = 1 nm, 1 nm data pitch, 2 s response, scan speed = 50 nm.min-1, averaged over 3 scans, zeroed at 350 nm). Samples were prepared in 100 μL (final volume) comprising 4 μM RNA in 10 mM lithium cacodylate buffer (pH 7.5), 1 mM KCl and 99 mM LiCl for spectra. RNAs were then heated at 70°C for 5 min and cooled down at room temperature for 1 h. For CD-melting, RNAs were dissolved at 4 μM in a solution containing 10 mM Li-cacodylate (pH 7.5) in 10 mM lithium cacodylate buffer (pH 7.2), and an increased K ^+^ concentration (1,10,100mM). Melting curves were obtained by recording the absorbance at fixed wavelengths (265 nm) as a function of temperature upon increasing the temperature from 20°C to 90°C at 0.2°C min-1. Final data were analyzed with Excel (Microsoft Corp.) and Prism

#### Fluorescence N-MethylMesoporphyrine (NMM) assay

Fluorescence assays were performed as previously described ^36^ The G4 folding RNAs or the control with a G/A mutant (200 pmol) were added to the folding buffer (20 mM Li-cacodylate (pH 7.5) and 100 mM LiCl or KCl). The reactions were then annealed to 70°C (5 min) and cooled down slowly at room temperature (1h). The appropriate buffer (20 mM Li-cacodylate (pH 7.5), 20 mM MgCl_2_, and 100 mM of either LiCl or KCl) was added to a final volume of 100 μL. Then, 2.5 eq/RNA of N-Methyl-Mesoporphyrin IX (NMM) (Frontier Scientific Inc., Logan, Utah) was then added and incubated for 5 min at room temperature (protected from light) in a 10 mm quartz cuvette. The fluorescence intensity was monitored using a Hitachi F-2500 fluorescence spectrophotometer with an excitation wavelength of 399 nm and the emission spectra were recorded between 500 nm and 650 nm. The fluorescence at 605 nm was used for quantification. All NMM assays were performed at least in duplicate.

#### RTS-stop assay

The reverse transcriptase stalling (RTS) assays were performed using in vitro synthesized RNA transcripts as described previously with the modifications described below ^35^. Each experiment was performed in the presence of 4.4 pmol of RNA and 5 pmol of Cy5-labeled primer in RTS buffer (50 mM Tris, 5 mM MgCl_2_, and 75 mM of either LiCl or KCl) in a final volume of 7.5 μL. A pairing step of 3 min at 75°C containing the RNA, the labeled primer and the RTS buffer without MgCl_2_ was performed. The temperature of the mixture was then decreased to 37 °C for 5 min and 0.5 μL of 0.1M DTT, 0.5 μL of 10 mM dNTPs and 1 μL of 25 mM MgCl_2_ were then added. Then, 1 μL of either 44 μM ligand (10 equiv/RNA) or water was added. In order to generate the ladders, 1 μL of 10 mM ddNTP (di-deoxyribonucleotide) was also incorporated into the reaction at this moment. Finally, 0.5 μL of purified MMuLV reverse transcriptase was then added to each tube and the reactions were incubated for 15 min at 45°C. The enzyme was then inactivated by adding 0.75 μL of 2N NaOH per sample and heating at 90°C for 10 min. Fifteen microliters of loading denaturing buffer (98% formamide, 10 mM EDTA) were added to each sample and they were then either stored at 4°C or were used for the denaturing gel migration. Ten microliters were loaded onto denaturing (8M Urea) 12% polyacrylamide gels (19:1) which were electrophoresed for 2 h at 50 W. The final images were generated using a GE Typhoon FLA 9000 fluorescence scanner (λex=635 nm), and are presented in Supplementary Figures.

### Pull-down assays via binding aptamers constructions

RBP enrichments was been performed as previously described ^47,48^. Cellular protein extracts were prepared from 4 confluent 15 cm dishes of U87 cells (3 for KCl and 1 for LiCl lysis buffer). The cell culture dishes were placed on ice and washed twice with ice-cold PBS (10 mL). The cells were then lysed by LiCl or KCl-lysis buffer (50 mM Tris-HCl (pH 7.5), 100 mM KCl or LiCL, 50 mM NaCl, 1 mM DTT, 0,1% Triton X-100, 10% glycerol, 2 mM MgCl_2_, 2 mM Vanadylribonucleosid complex RNase inhibitor (New England Biolabs) and mini Complete Protease inhibitors (Roche) for 30 min at 4°C. Debris from the lysate was then removed by centrifugation at 17 000 x g for 20 min at 4°C, resulting in a supernatant of 4000uL. The solution was then concentrated using Amicon Ultra-4 MWCO 3 000 centrifuge tubes (EMD Millipore) at 1 700 x g until the final volume required 500 uL/ sample. The protein concentrations in the extract were then determined by Bradfords method according to BIO-RAD instructions. The extracts were then pre-cleared by the addition of 50 µL of a 50% slurry of Streptavidin Sepharose High Performance beads (GE Healthcare) and incubated for 16 h at 4°C. The beads were discarded, and the pre-cleared lysate was supplemented with 1.5uL of RNasin and kept for RNA-coupled Sephrasose beads step.For the RNA coupling step : the in vitro transcribed 4xS1m control, 4xS1m WT or 4xS1m G/A mutant RNA (30 µg) was dissolved in 140 µL of KCl-lysis buffer without MgCl_2_ or RNase inhibitors and then heated to 70°C and slow cooled at rate of 1*C/min until 1 hour. The streptavidin beads (100 µL) were washed three times with appropriate lysis buffer and the RNA transcript. Subsequently, the solution was supplemented to a final concentrations of 1 mM MgCl_2_ and 120 U of RNase inhibitors were added (Protein purification platform from Université de Sherbrooke). The RNA solutions were then mixed with 100 µL of a pre-washed (with KCl-lysis buffer) Streptavidin Sepharose bead slurry and incubated for 2 h at 4°C. After this incubation, the beads were centrifuged at 20 x g for 1 min and the supernatant was discarded. The beads were then washed once with ice-cold KCl-lysis buffer. Next, the pre-cleared lysate (e.g. the one incubated 16 h at 4°C) was supplemented with 1.5 µL of RNase inhibitor (Protein purification platform from Université de Sherbrooke), added to the prepared RNA-coupled Streptavidin Sepharose beads and incubated for 3 h at 4°C. Lastly, the beads were washed five times with 1 mL of KCl-wash buffer (50 mM Tris-HCl (pH 7.5), 100 mM KCl, 200 mM NaCl, 1 mM DTT, 0,1% Triton X-100, 10% glycerol, 5 mM MgCl_2_, Ribonucleoside Vanadyl complex RNase inhibitors (NEB) and Mini Complete Protease inhibitors (Roche)).

### Mass spectrometry

Sample preparation included the reduction and the alkylation of the proteins followed by trypsin digestion. Digested peptides were then purified on C18 columns. All these steps were performed as described previously 49.

The raw files from LC-MS/MS were analyzed using the MaxQuant version 1.6.2.2 software and the Uniprot human database. MaxQuant supports the label-free quantification of protein. The settings used for the MaxQuant analysis were: i) 2 miscleavages were allowed; ii) enzymes were Trypsin (K/R not before P); and, iii) variable modifications included in the analysis were methionine oxidation and protein N-terminal acetylation. Mass tolerances of 7 ppm for precursor ions and 20 ppm for fragment ionswere used. In addition, the following parameters were used: label-free quantification with an LFQ minimum ratio count of 2; identification values “PSM FDR”; “Protein FDR” and “Site decoy fraction” 0.05; and, “Match between runs”. Proteins positive for at least one of the “Reverse” and “Potential contaminant” categories were eliminated. MS/MS count were used to determine the fold enrichment in WT K+ vs Li+ samples for each protein identified. Proteins with a coverage >5 %, a fold enrichment of >2.0 and a MaxQuant score >15 were conserved for subsequent analysis.

